# TraIT, Translational research IT bridging the valley of death in translational medicine

**DOI:** 10.1101/2020.05.18.052563

**Authors:** Jan-Willem Boiten, Rita Azevedo, Kees van Bochove, Marinel Cavelaars, André Dekker, Remond J.A. Fijneman, Peter Lansberg, Wim van der Linden, Barend Mons, Nikolas Stathonikos, Henk M.W. Verheul, Jeroen A.M. Beliën, Gerrit A. Meijer

## Abstract

Translating new technology and biological findings into clinical applications is hampered by insufficient translational research IT. The Dutch Translational research IT (TraIT) initiative organizes, deploys, and manages data and workflows in an on-line “office suite”, supplemented with efficient training and user support. TraIT has been adopted by a wide user community providing an excellent large-scale demonstrator for the nation-wide Health-RI initiative.

## Introduction

Tremendous progress in biology and technology, exemplified by the “omics” revolution, heralds a bright future for healthcare. Yet, translating these discoveries into new diagnostics, drugs and health technologies faces severe challenges, which slows their implementation in daily patient care. Intriguingly, this phenomenon, often referred to as the “translational valley of death” ^[1]^, is not so much caused by factors in the domains of biology and “omics” themselves, but rather in the lower-profile areas of organization, logistics, and data management. In other parts of society, exactly these domains have largely benefited from information technology (IT). There is every reason to believe that elevating IT in biomedical translational research to the level we encounter when shopping on the internet, or managing our personal finances online, can largely bridge this valley.

### The translational paradigm

The mission of biomedical research is to produce better outcomes for patients, as well as improving health through new preventive measures and interventions. The central paradigm is that a better understanding of how disease biology affects disease phenotype, i.e. clinical presentation and outcome, will enable disease prevention or treatment with better outcomes. This translational paradigm is valid across many health and disease areas, but also across different types of (unmet) clinical need such as prevention, early detection, therapeutic intervention and response monitoring.

### Evidence-based medicine

The principles of evidence-based medicine come into play when we aim at improving health outcomes by translational research. In order to improve outcomes, we need to change clinical practice and practice guidelines, as well as provide the evidence to support such changes. Currently, however, only a fraction of the many translational research studies in medicine yield practice-changing evidence for various reasons. ^[2]^ To increase the speed at which we bring basic research innovations to patients, we need to improve operations, logistics and data management of studies that produce the required evidence.

Previously, many initiatives in this field have been isolated efforts focusing merely on parts of the problem without considering the full process they are supposed to facilitate. Interestingly, while in our daily office work we all are used to integrated IT solutions, scientists are still lacking such a “translational research office suite”.

### Data generation and integration

Data integration initiatives are at risk of being monolithic, often starting halfway down the data generation process and assuming that data arise spontaneously. Showcase initiatives in this domain had to invest into time-consuming, costly, and often manual efforts to interactively polish data in order to make these interoperable, an underexposed activity referred to as data curation or ETL (extract-transform-load). ^[3,4]^ Basically, all of this is manual labor needed to compensate for inefficient organization of the research process. Moreover, in many data integration initiatives active translational and/or clinical researchers are underrepresented.

The field of “big data” in translational research is challenging. The challenge is not in capturing, storing, processing and analyzing petabytes of data produced by next-generation sequencing (NGS) and imaging. Most technical challenges in these domains have been or are being addressed. The real issues here are related to data access. Because success in translational research depends on finding meaningful associations between omics and phenotypic data, the availability of high-quality, high-volume clinical phenotype data is as important as the quality of the omics data. For the latter, we rely on standard operating procedures in the lab and thorough metadata registration when biobanking the samples. For clinical phenotype data in most instances we turn to data from hospital records that are often still unstructured, of different granularity, spread across multiple (often not integrated/linked) systems, mostly recorded in the local language, and inherently more subjective than the biological data. So, access to high-quality data is the weak link in the translational research process chain.

### Biomarker validation

Most biomarker studies never achieve the level of sophistication of practice-changing clinical intervention trials. In most cases, after a case-control proof-of-concept study, only a retrospective validation study is provided, although in general this is considered insufficient to change clinical practice. Unfortunately, it stops here for most candidate biomarkers. One of the likely reasons why practice-changing prospective validation is rarely undertaken, is because it is considered being too complex. In addition, studies on the same biomarker by different institutions often yield conflicting results, which relates at least in part to data issues. It is evident the challenges in data reproducibility in translational research extend beyond preclinical drug research ^[5]^, and also affect e.g. biomarker research. ^[6,7]^ Opportunities to escape from this deadlock are to increase the quality of retrospective data for clinical and translational research and to facilitate solid prospective biomarker validation studies. Both can be largely facilitated by an IT infrastructure that supports close interactions between research and care workflows.

## CTMM-TraIT

Against this background, the CTMM-TraIT (http://trait.health-ri.nl) initiative was launched in the Netherlands. The Center for Translational Molecular Medicine (CTMM; http://lygature.org/ctmm-portfolio) was a €300 million public-private partnership focusing on molecular imaging and laboratory diagnostics in oncology and cardiovascular, neurodegenerative and inflammatory diseases. TraIT (Translational Research IT) was a national consortium that organizes, deploys, and manages a multi-partner IT infrastructure for data and workflow management, targeting biomarker research. TraIT enables integrating and interrogating information across the four major domains of translational research: clinical, imaging, biobanking and experimental or molecular (omics) data. The TraIT project was launched late 2011, and has gained national and international recognition since.

### The TraIT approach

TraIT aims to facilitate the standard workflow of translational studies that address specific unmet clinical needs. At the input side, patients enter the research domain; at the output side, the primary goal is to improve healthcare through large-scale data integration generating new knowledge. In addition, scientific publications, intellectual property, improved healthcare and, increasingly important, opportunities to interact with the public are aimed for as well (Figure 1).

TraIT is workflow-oriented and driven by the needs and perspectives of translational/clinical researchers. Furthermore, TraIT avoids duplication of efforts by adopting (and adapting), wherever possible, existing standards, technologies and operational applications.

**Figure 1.**
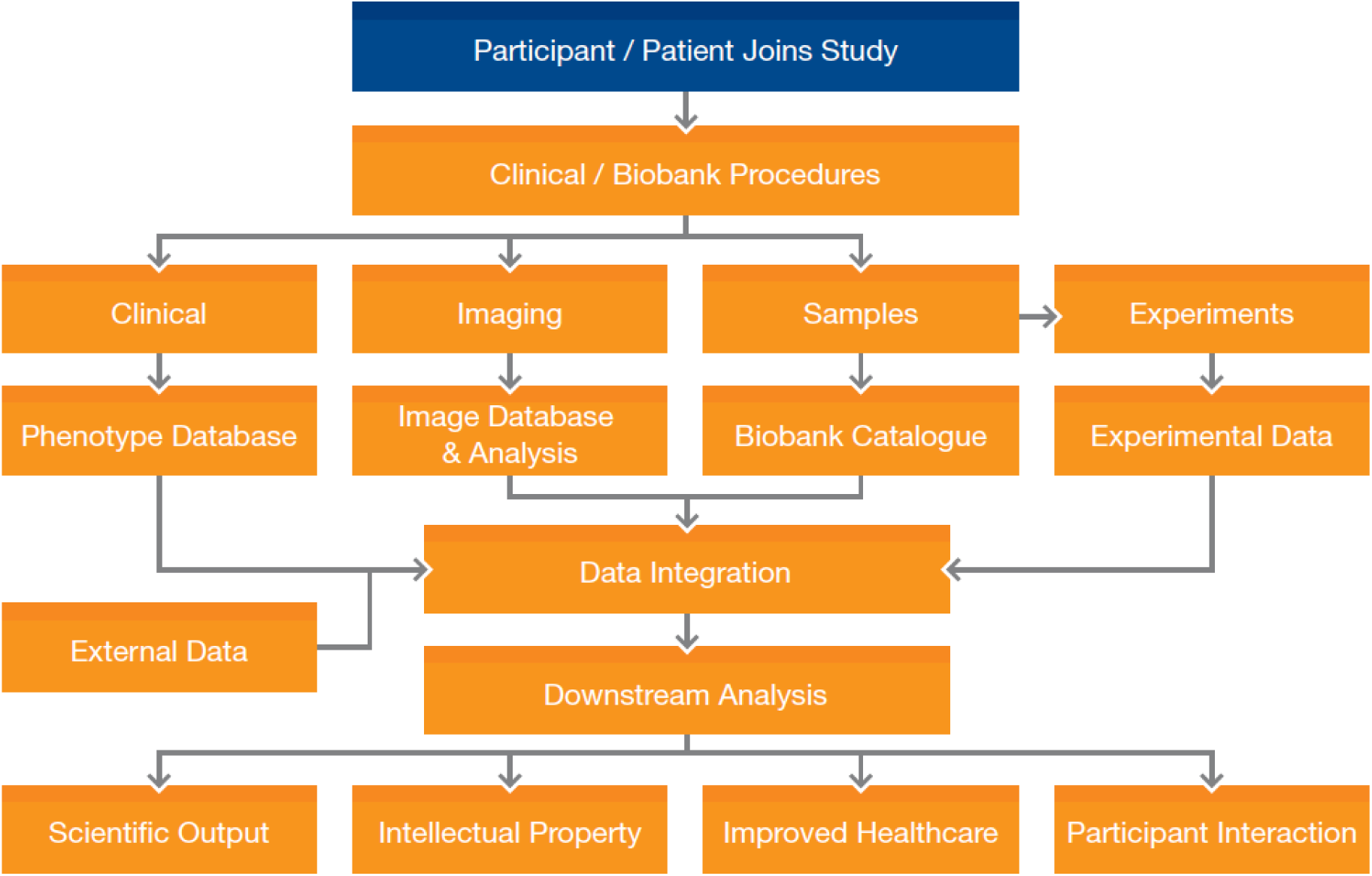
The standard translational research workflow supported by TraIT. An individual enters a study as a patient in a clinic or a donor to a biobank. In a series of standard procedures, data are generated and stored in the phenotype annotation, imaging, biobanking and experimental domains, and used in downstream analysis to achieve better health outcomes for future patients.

### Care and research data integration

The crucial step is to integrate phenotype and omics data to identify those associations across domains, which may lay the basis for new diagnostic or therapeutic applications. This process of data integration nowadays increasingly involves integration with publicly available datasets. In line with the principle of avoiding duplication of efforts, TraIT has adopted existing successful applications, often open source, to fill crucial slots in its workflow. For clinical data capture, the open-source application OpenClinica has been selected as a tool. (https://www.openclinica.com/) It has appropriate quality assurance features including an audit trail.

The Extensible Neuroimaging Archive Toolkit (XNAT; https://www.xnat.org) is used as the tool for clinical image collection and sharing, while it also provides resources for sharing image analysis results. Next to clinical imaging, digital pathology was supported through the tEPIS (https://trait.health-ri.nl/trait-tools/tepis/) system based on the Philips Digital Pathology solution.

In the biobanking domain, biobank catalogues of different dimensions, ranging from individual studies up to the Dutch national pathology registry PALGA with >50 million entries (http://www.palgaopenbaredatabank.nl/), feed into the TraIT workflow, and can be interrogated from TraIT. In this area TraIT has adopted the “catalogue” concept developed by BBMRI-NL, the Dutch node in the European biobanking network. (https://catalogue.bbmri.nl/)

A major challenge in the experimental data domain is to establish standardized pipelines, avoiding IT complexity in the transfer of methods from the bioinformatics expert to the broader medical research community. By using Galaxy, such mature workflows for genomics and proteomics data processing can be made available to non-bioinformatician translational researchers. (https://galaxyproject.org)

Data integration is the centerpiece of translational research IT. After careful evaluation of available options, TraIT has adopted the open-source solution tranSMART for this purpose in late 2012. (http://transmartfoundation.org/) tranSMART aims to provide “hypothesis-free” data browsing of the combined clinical and genomics data set of biomarker studies. This allows addressing research questions on the spot, e.g. to determine the impact of certain gene mutations on the responsiveness of colon cancer to a particular systemic therapy. After the decision to join the tranSMART community was taken, TraIT has contributed significantly to the further development of the tranSMART solution, in particular driven by two showcase biomarker projects in colon cancer and prostate cancer. The TraIT enhancements have become an integral part of the community release of tranSMART.

## What has been achieved?

TraIT has implemented the complete suite of tools needed to support the personalized medicine research workflow (Figure 1) and makes this available online (https://trait.health-ri.nl/trait-tools). On top of this, users are supported by a helpdesk, documentation, training, and a self-service portal, addressing most of the obvious questions by novice users, as well as account management for more complex research programs. This additional layer of services has been key to the success of TraIT.

Table 1 illustrates penetration of TraIT in biomedical research, mostly in The Netherlands but also across research institutes abroad that participate in multicenter studies. It shows use over a wide range of disease areas and disciplines, albeit with a clear predominance of oncology. In total, TraIT has about 4250 unique users from The Netherlands and abroad, who apply TraIT tools in approximately 420 studies. Complex studies running at multiple sites and in multiple research domains (clinical, imaging, etc.) potentially benefit most from the closely integrated services of TraIT.

**Table 1.** Number of studies using TraIT by medical discipline and breadth of use (2011-2019).

| Discipline | Single site, single tool | Single site, multitool | Multisite, single tool | Multisite, multitool | Total |
| --- | --- | --- | --- | --- | --- |
| Oncology | 59 | 8 | 98 | 23 | 188 |
| Cardiovascular | 18 |  | 29 | 4 | 51 |
| Neurology | 17 |  | 21 | 4 | 42 |
| Internal medicine | 17 |  | 23 | 1 | 41 |
| Immunology | 11 |  | 12 | 2 | 25 |
| Gynaecology | 3 |  | 16 |  | 19 |
| Surgery | 5 | 1 | 12 |  | 18 |
| Other | 17 | 1 | 18 | 1 | 37 |
| **TOTAL** | **147** | **10** | **229** | **35** | **421** |

## Discussion

TraIT is a user-driven, process-oriented translational research IT initiative, aiming to facilitate data stewardship of translational research projects from start to end. Current fragmented approaches in this field often are a critical hurdle in taking biomarker proof of concepts to the next level of validation and ultimately patient care.

### Technical aspects

Making the workflow presented in Figure 1 operational in a fully automated manner requires optimal syntactic and semantic interoperability between the respective applications. Semantic interoperability is an important hurdle, especially in the clinical domain, but also when it comes to image annotations. Much of the clinical information is still unstructured text in electronic hospital records. Structured or synoptic reporting is beginning to gain ground in clinical reporting, but is still in its infancy.^[8]^

In addition, high-quality data stewardship and re-use of research data are increasingly gaining attention, e.g. through the GO FAIR initiative (www.go-fair.org), in which FAIR stands for “findable, accessible, interoperable and reusable” data and services.^[9]^ To achieve this goal, not only IT systems like TraIT should comply with community emerging standards, but also researchers. While this is easier said than done, it can be largely facilitated by an application suite like TraIT enabling “compliance by design”.

### Budgetary aspects

It is evident that a comprehensive translational research IT infrastructure requires substantial budget. TraIT initially had a budget of €16 million for four years. TraIT has restricted spending on custom software development, the traditional glutton in many IT projects, by relying on existing solutions. Instead, building user communities, bottom-up good practice development, and subsequent deployment and professional-level user support were prominent budget items in TraIT.

The preference for open-source solutions facilitates community-based collaboration, like on the data integration application tranSMART. In this way substantial efforts from both academic and pharmaceutical partners were made available for a joint effort, resulting in separate development branches of tranSMART having been merged into one. Sustaining these community efforts over many years proves to be challenging though.

We believe that TraIT is not the final answer to all translational research data stewardship challenges, but rather a start. While the TraIT project had a substantial budget, this was merely enough to set the flywheel in motion. The pioneering activities of TraIT have provided a (large-scale) proof-of-concept demonstrating that a well-coordinated bottom-up approach could offer robust data infrastructure services. Among stakeholders there is increasing awareness of the problem of data access and logistics as well as willingness to act in a concerted way. In the Netherlands academia, medical centers, funders, patients, private partners, and government have joined forces to establish a comprehensive national infrastructure for personalized medicine and health research, called Health-RI (https://health-ri.nl), which we envision to be the future sustainable landing zone for TraIT. The Dutch Cancer Society has supported this strategy with a € 2.25 million grant to transition TraIT into Health-RI.

### Scientific community

Translational research in biomedicine is a complex process that requires a dedicated infrastructure and excellent teamwork, much like running an industrial process.

This conflicts with traditional academic structures and career models based on individual excellence and originality. Current academic reward systems merely recognize prominent authorship, rather than contributions to the process chain of translational research. ^[10]^ In fact, this is a major obstacle to data sharing, hampering value creation for patients. While this is true for translational research in general, it also applies to IT initiatives and bioinformatics. When requests for funding proposals are issued, these often call for innovative cutting-edge initiatives, rather than for consolidating and extending existing successful infrastructures.

### The public at large

Biomedical research traditionally receives substantial public attention, often through media coverage heralding the clinical impact of findings that are hardly past the discovery stage. However, interactions with the public have changed from a sending/receiving mode to participatory. Public engagement comes in different flavors, ranging from patient advocacy, advice on matters of ethical, legal, and societal implications, to demanding more control over one’s personal data.

### Public-private partnerships

Some initiatives, like CTMM at the Dutch, and EATRIS (https://eatris.eu) at the European level, address translational research more explicitly than others. CTMM and EATRIS arose from the notion that successful implementation of the products of translational research, e.g. new drugs, diagnostics, or therapeutic interventions, is usually accomplished in partnerships with well-established private entities. Any IT solution supporting translational research should therefore be optimally positioned to support public-private partnerships.

Within TraIT we took this concept one step further: the TraIT consortium itself has been set up as a public-private consortium involving commercial organizations developing, producing and marketing biomedical software and pharmaceutical and medical technologies. This concept has proven to be very productive.

## Future perspectives

Initiatives on research infrastructure like the European Strategy Forum on Research Infrastructures (ESFRI), funded by the European Union and national governments, are important steps forward. At the same time, they are dispersed and domain-rather than process-oriented. Internet became a success through the adoption of simple standards and browsers that allowed maximum freedom to implement and to meet the needs of many end-users. The GO FAIR initiative, following the recommendations for the European Open Science Cloud as an ‘Internet of FAIR data and Services, and in the health field for instance the Global Alliance for Genomics and Health (http://genomicsandhealth.org/), are pursuing this goal. Health-RI in the Netherlands is also committed to this approach. Similarly, research infrastructures may need to join forces to facilitate end-users in the universal process of translational research. Integrated IT solutions for translational research, like TraIT’s “office suite” approach, are crucial for translational research to deliver on its expectations. The TraIT project has been pioneering in this field and as such did form an important foundation for the Health-RI initiative.

